# Expression Profiles of FABP4 and FABP5 in Breast Cancer: Clinical Implications and Perspectives

**DOI:** 10.1101/2024.10.01.616073

**Authors:** Xingshan Jiang, Yiqin Xiong, Jianyu Yu, Anthony Avellino, Shanshan Liu, Xiaochun Han, Zhaohua Wang, Jonathan S. Shilyansky, Melissa A. Curry, Jiaqing Hao, Edward R. Sauter, Sonia L. Sugg, Bing Li

**Affiliations:** Department of Pathology, University of Iowa, Iowa City, IA, USA; Holden Comprehensive Cancer Center, University of Iowa, Iowa City, IA, USA; Division of Cancer Prevention, NIH/NCI, Rockville, MD, USA; Department of Surgery, University of Iowa, Iowa City, IA, USA

**Author notes:** **Corresponding Author** Bing Li, Professor at Department of Pathology, University of Iowa. Address: 431 Newton Road, Iowa City, IA, 52242, USA,.

## Abstract

The incidence of breast cancer continues to rise each year despite significant advances in diagnosis and treatment. Obesity-associated dysregulated lipid metabolism is believed to contribute to the increasing risk of breast cancer. However, the mechanisms linking lipid dysregulation to breast cancer risk and progression remain to be determined. The family of fatty acid binding proteins (FABPs) evolves to facilitate lipid transport and metabolism. As the predominant isoforms of FABP members expressed in breast tissue, adipose FABP (A-FABP, also known as FABP4) and epithelial FABP (E-FABP, FABP5) have been shown to play critical roles in breast carcinogenesis. In this study, we collected surgical breast tissue samples from 96 women with different subtypes of breast cancer and comprehensively analyzed the expression pattens of FABP4 and FABP5. We found that distinct expression profiles of FABP4 and FABP5 were associated with their unique roles in breast cancer development. FABP4, mainly expressed in breast stroma, especially in adipose tissue, supported neighboring tumor cell lymphvascular invasion through secretion from adipocytes. In contrast, FABP5, primarily expressed in epithelial-derived tumor cells, promoted tumor metastasis by enhancing lipid metabolism. Thus, elevated levels of FABP4 and FABP5 could serve as poor prognostic markers for breast cancer. Inhibiting the activity of FABP4 and/or FABP5 may offer a novel strategy for breast cancer treatment.

## Introduction

Despite advancements in diagnosis and treatment, breast cancer remains a leading cause of death among women in the U.S. and worldwide^1^. The global incidence has surged from 641,000 cases in 1980 to over 2.3 million in 2020^1, 2^. While traditional risk factors such as aging, genetic mutations, family history, and reproductive status play a significant role, obesity has emerged as an additional contributor to the risk of various cancers, including breast cancer^3-5^. However, the molecular mechanisms that link obesity to increased breast cancer risk remain largely unexplained.

Obesity is characterized by dysregulated lipid metabolism, leading to excess lipid accumulation in various organs, tissues, and cells^6^. Given the insoluble nature of lipids in the human body, a family of proteins known as fatty acid binding proteins (FABPs) has evolved to solubilize fatty acids, facilitating their transport and metabolism^7, 8^. The FABP family consists of at least nine members, each with a characteristic tissue distribution, such as liver FABP (L-FABP, FABP1), intestinal FABP (I-FABP, FABP2), heart FABP (H-FABPT, FABP3), adipose FABP (A-FABP, FABP4), epithelial FABP (E-FABP, FABP5), etc^9, 10^. Since breast tissue primarily consists of adipose tissue, epithelial ducts and lobules, FABP4 and FAPB5 are the predominant members involved in coordinating lipid metabolism and maintaining lipid homeostasis in breast tissue^11^. It is of great interest to understand whether and how FABP4 and FAPB5 link dysregulated lipid metabolism to breast cancer risk and progression.

Emerging evidence from our group and others has shown that FABP4 and FABP5 play critical roles in breast cancer development and progression in preclinical models^10, 12-15^. For example, obesity induced by a high-fat diet increases the secretion of FABP4 from adipocytes and macrophages into the circulation where soluble FABP4 can directly target breast cancer, inducing ALDH1-mediated breast tumor stemness and invasiveness^16, 17^. In contrast, FABP5 expression in estrogen receptor (ER)-negative cancer cells was shown to promote tumor growth and is associated with a poor prognosis^10, 15^. These findings suggest that FABP4 and FABP5 could serve as important prognostic markers for breast cancer. In this study, we comprehensively analyzed the expression profiles of FABP4 and FABP5 in both normal and breast cancer tissues to assess their clinical significance and potential therapeutic implications.

### Expression profiles of FABP4 and FABP5 in normal adjacent breast tissue

To investigate the role of FABP4 and FABP5 in breast cancer development, we first assessed their expression patterns in normal adjacent breast tissue specimens collected from patients with breast cancer. Normal adjacent breast tissue primarily consists of lobules (Figure 1A), ducts (Figure 1B), adipose tissue (Figure 1C), blood vessels (Figure 1D) and interlobular connective tissue, which provides structural support to the breast and helps maintain its function^18^. Histologically, lobules are mainly composed of milk-producing luminal epithelial cells (red arrows in Figure 1A) and an outer layer of contractile myoepithelial cells (black arrows in Figure 1A), which help propel milk from the lobules into the ducts. Similar to lobules, ducts are formed by luminal epithelial cells and myoepithelial cells, serving as a passageway for milk flow from the lobules toward the nipple. Immune cells, such as mononuclear macrophages, are present in both lobules and ducts, which respond to potential invaders and maintain tissue homeostasis (green arrows in Figure 1A and Figure 1B).

**Fig. 1.**
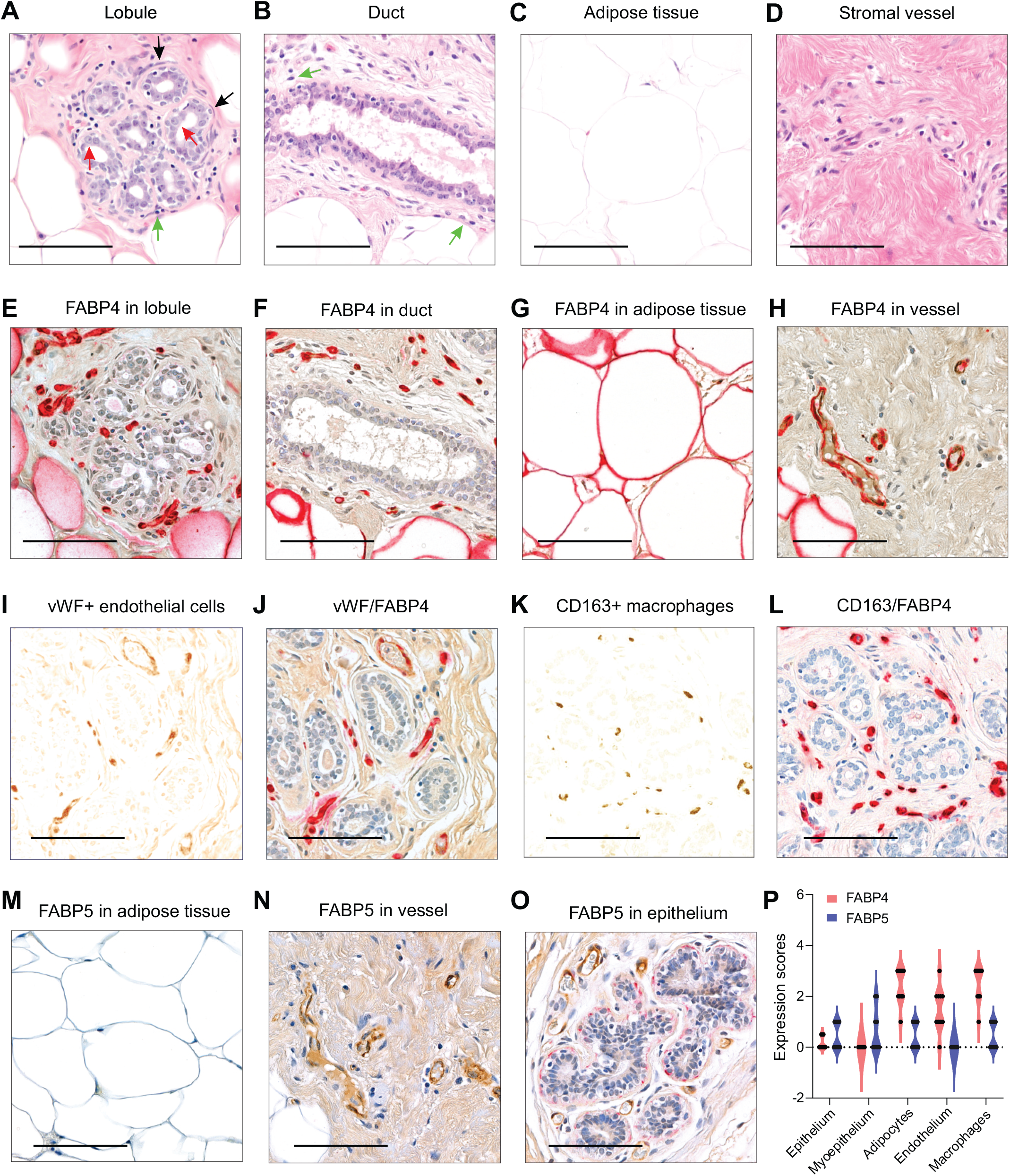
Expression profiles of FABP4 and FABP5 in normal breast tissue. (**A-D**) Representative hematoxylin and eosin (H&E) staining of normal breast tissue, showing lobules (**A**), ducts (**B**), adipocytes (**C**), and blood vessels (**D**). Scale bar: 100 μm. (**E-H**) Representative immunostaining for FABP4 (red) in normal breast tissue, including lobules (**E**), ducts (**F**), adipocytes (**G**), and blood vessels (**H)**. Scale bar: 100 μm. **(I, J**). Analysis of colocalization of vWF (brown) (**I**) and FABP4 (red) (**J**) expression in normal breast tissue. Scale bar: 100 μm. (**K, L**) Analysis of colocalization of CD163 (brown) (**K**) and FABP4 (red) (**L**) expression in normal breast tissue. Scale bar: 100 μm. (**M-O**) Representative immunostaining for FABP5 (red) and vWF (brown) in normal breast tissue, showing adipocytes (**M**), blood vessels (**N**), and epithelium (**O**). Scale bar: 100 μm. (**P**) Comparison of FABP4 and FABP5 expression levels across different types of cells in normal breast, including epithelium, myoepithelium, adipocytes, endothelium, and macrophages (n=9 for each group)

Immunohistochemical (IHC) staining showed that FABP4 was absent in both luminal epithelial and myoepithelial cells (Figure 1E, 1F). Instead, it was highly expressed in the breast stroma, including adipocytes (Figure 1G) and vessels (Figure 1H). Using endothelial marker von Willebrand factor (vWF) (Figure 1I) and macrophage marker CD163 (Figure 1K), we further demonstrated that FABP4 was highly expressed in vWF^+^ endothelial cells (Figure 1J) and CD163^+^ macrophages (Figure 1L). Unlike the stromal cell-specific expression pattern of FABP4, FABP5 was either absent or weakly expressed in the stroma of normal breast tissue, including adipose tissue (Figure 1M) and vessel endothelium (Figure 1N). However, FABP5 expression was observed in myoepithelial and epithelial cells in half of the samples (Figure 1O), suggesting a specific role of FABP5 in epithelial cell function. Given the different expression patterns of FABP4 and FABP5 (Figure 1P), it is likely that FABP4 plays a more prominent role in supporting lipid metabolism and functions in stromal cells, while FABP5 may be involved in epithelial cell activity in normal breast tissue.

### FABP4 expression pattern in breast cancer tissue

To investigate the role of FABP4 in breast cancer development and progression, we collected surgical breast cancer tissues from 96 women diagnosed with various subtypes of breast cancer (Table 1) and analyzed FABP4 expression patterns in these samples. In patients diagnosed with ductal carcinoma in situ (DCIS), cancer cells were confined within the milk ducts (Figure 2A) without spreading into surrounding stromal tissue, including immune cells (Figure 2B), adipose tissue (Figure 2C) and vessels (Figure 2D). FABP4 expression exhibited a similar pattern to that in normal breast tissue: absent in epithelial cells (Figure 2E), but high in CD163+ macrophages (Figure 2F), adipocytes (Figure 2G), and vessels (Figure 2H). In patients with invasive breast cancer, most tumor cells were FABP4 negative (Figure 2I, 2J), though some tumor cells, particularly those near to adipose tissue, exhibited positive staining for FABP4 (Figure 2K). Notably, the closer the tumor cells were to adipose tissue, the stronger the FABP4 staining in these cells, suggesting that the ectopic FABP4 staining in these invasive frontline tumor cells may originate from adipose tissue. Indeed, we noticed that the size of adipocytes close to the tumors was significantly smaller than those away from tumors (Figure 2L-2N). Previous findings from our group and others demonstrated that FABP4 secreted by adipose tissue could serve as an adipokine, providing energy support and promoting oncogenic signaling activation in tumor cells^14, 17, 19^. Our current findings provide further evidence that tumor cells could be taking up adipose tissue derived FABP4 for their invasive benefits at the frontline.

**Fig. 2.**
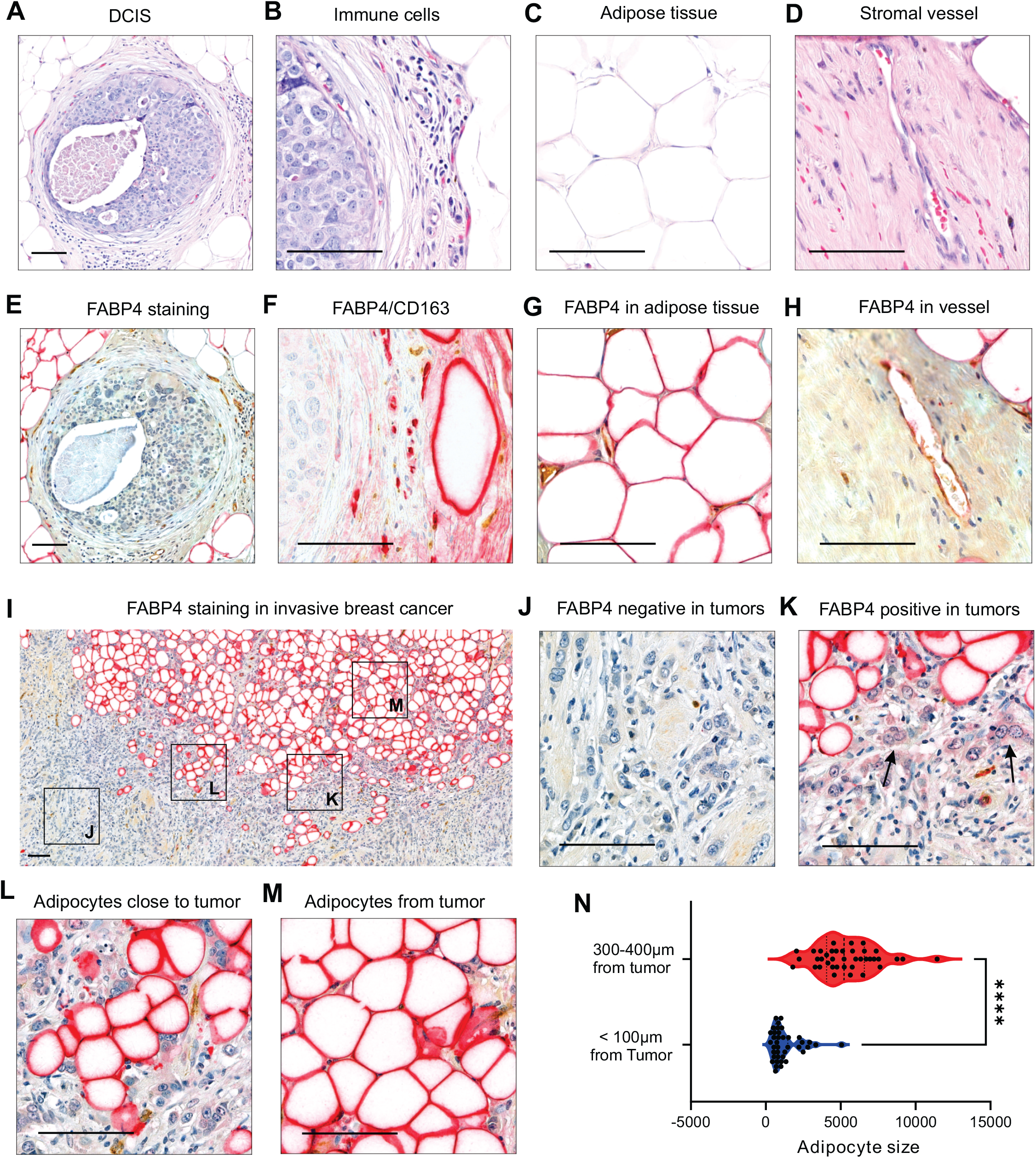
FABP4 expression pattern in breast cancer tissue. (**A-D**) Representative H&E staining of ductal carcinoma in situ (DCIS), including DCIS regions (**A**), surrounding immune cells (**B**), adipocytes (**C**), and stromal vessel (**D**). Scale bars: 100 μm. (**E-J**) Representative immunostaining for FABP4 (red), CD163 (brown) and VWF (brown) in DCIS breast tissue, including DCIS regions (**E**), immune cells (**F**), adipocytes (**G**), and stromal vessel (**H**). Scale bars: 100 μm. (**I-M**). Representative immunostaining for FABP4 in invasive frontline tumor cells and adjacent adipose tissue. Most tumor cells were negative for FABP4(J). Tumor cells close to adipocytes were positive for FABP4 (black arrow) (K). Adipocytes adjacent to tumor cells were small (L). Adipocytes away from tumor were large (M). Scale bar for I: 200 μm, scale bar for panel J-M: 100 μm, (**N**) Comparison of adipocyte sizes at varying distances from tumor cells (p<0.0001, n=40 cells).

### FABP5 expression pattern in breast cancer tissue

Next, we analyzed FABP5 expression pattern in the same cohort. Similar to our observations in normal breast tissue, DCIS tissues showed weak FABP5 expression in epithelial tumor cells and myoepithelial cells (Figure 3A), CD163+ macrophages (Figure 3B), adipocytes (Figure 3C) and vessels (Figure 3D). However, invasive breast cancer tissues exhibited a significant increase in FABP5 expression (Figure 3E), mainly in tumor cells (Figure 3F). Vessel endothelial cells also exhibited increased expression of FABP5 (Figure 3G). In contrast to FABP4, which was primarily expressed in tumors close to adipose tissue (Figure 2K), the pattern of FABP5 expression in tumor cells was either ubiquitously (Figure 3E) or clustered in groups of cells sporadically distributed throughout the tumor (Figure 3H). Statistical correlation analysis showed that there were no correlations between the expression pattern of FABP4 and FABP5 in tumor cells (Figure 3I). In addition, in tumor cells overall expression of FABP5 was significantly higher than FABP4 (Figure 3J). Altogether, the different expression patterns/levels of FABP4 and FAPB5 in invasive breast cancer suggested their distinct roles in breast cancer development and progression.

**Fig. 3.**
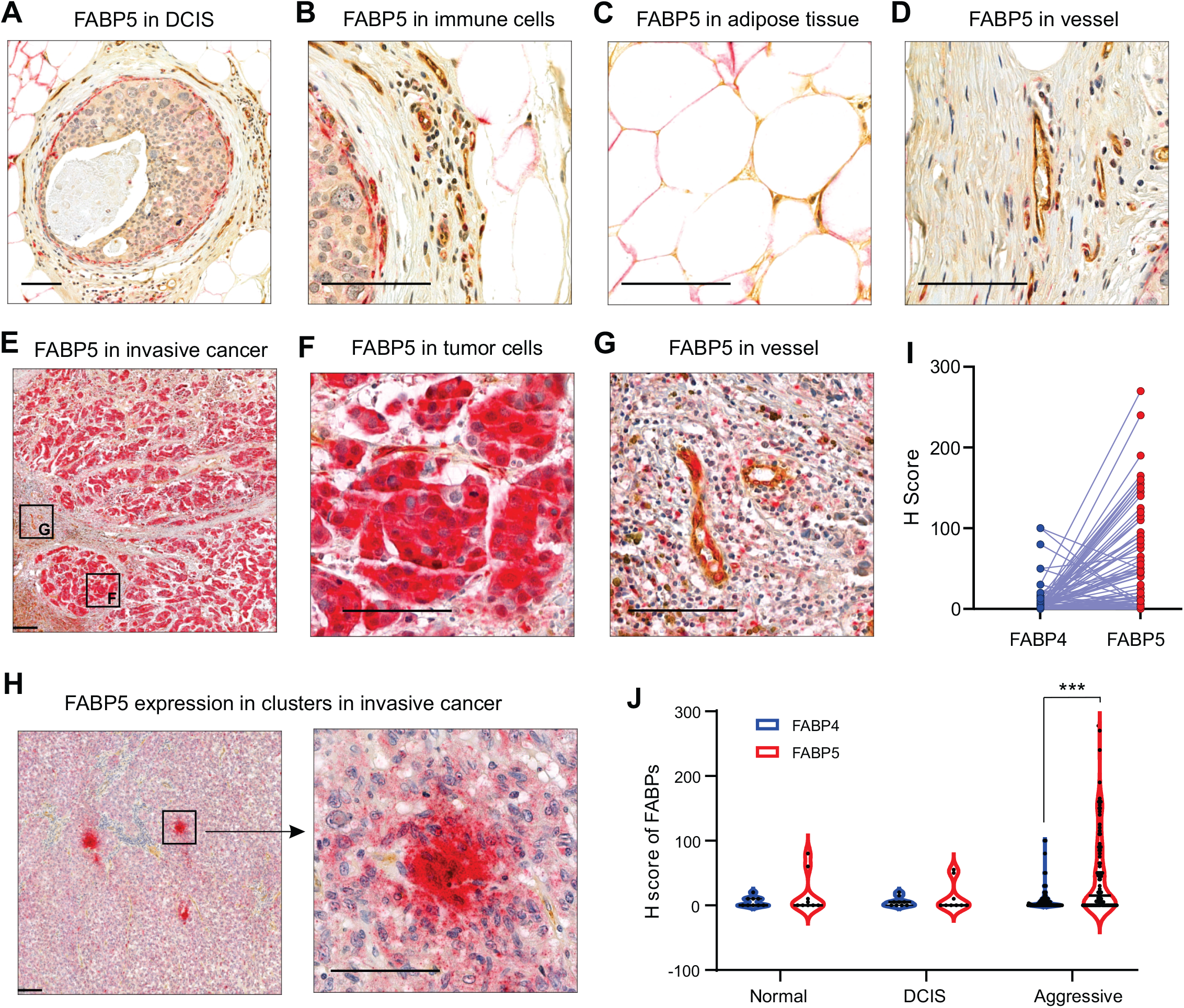
FABP5 expression pattern in breast cancer tissue. (**A-D**) Representative immunostaining for FABP5 (red) and vWF (brown) in DCIS, including DCIS regions (**A**), surrounding immune cells (**B**), adipocytes (**C**), and stromal vessel (**D**). Scale bars: 100 μm. (**E-H**) Representative immunostaining for FABP5 (red) and vWF (brown) in invasive breast cancer, showing two distinct patterns of FABP5 expression in tumor cells: ubiquitous expression in tumor cells (**E, F**) or clusters of positive cells sporadically distributed throughout the tumor tissue (**H**). Scale bars: 100 μm. (**I**) Correlation analysis between FABP4 and FABP5 expression levels in tumor cells of patients with breast cancer (n=96). (**J**) Comparison of FABP4 and FABP5 expression levels across normal breast tissue, DCIS, and aggressive cancer (p<0.001, n=96).

### FABP4 expression at the tumor border correlates with lymphovascular invasion

To dissect the potential role of FABP4 in breast cancer development and progression, we first analyzed the correlation of FABP4 expression in tumor cells with various clinical parameters, including age (Figure 4A), BMI (Figure 4B), tumor size (Figure 4C), Elston-Ellis grade (Figure 4D), metastasis (Figure 4E), and patient vital status (Figure 4F). None of these clinical parameters showed a significant correlation with FABP4 expression in tumors except the Elston-Ellis grade (Figure 4D), which showed a weak but significant correlation with FABP4 H score in tumor cells. We further assessed FABP4 expression across different breast cancer subtypes, including hormone sensitive (ER+/PR+/HER2-), HER2 positive (ER-/PR-/HER2+), and triple negative (ER-/PR-/HER2-). The overall expression levels of FABP4 in tumor cells were very low across breast cancer subtypes, and there were no significant differences in FABP4 expression H score among various subtypes (Figure 4G), suggesting minimal impact of hormones (*e*.*g*. estrogens, progesterone) or growth factor signals (HER2) on FABP4 expression. As FABP4-positive tumor cells were mainly located in regions that often interacted with adipose tissue (Figure 2K), we further determined if FABP4 expression at the tumor/adipose tissue border (Figure 4H, 4I) impacted clinical variables. Interestingly, FABP4 expression at the tumor border was positively correlated with tumor lymphovascular invasion (Figure 4J). Given tumor cells mainly spread through the lymphatic and vascular systems^20^, FABP4 may mediate adipose/tumor crosstalk by enhancing tumor invasion.

**Fig. 4.**
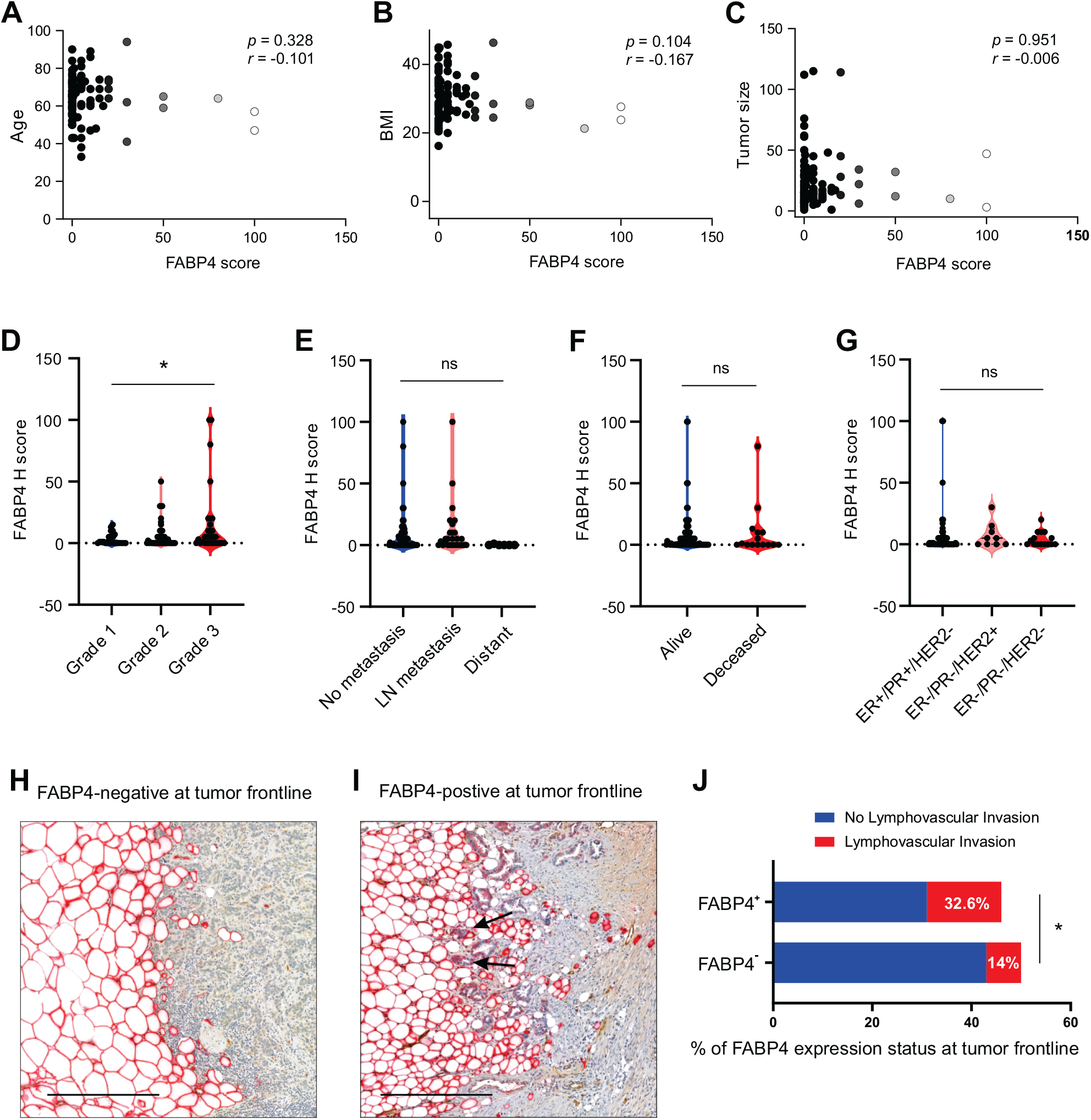
Correlations of FABP4 expression in tumor cells with clinical parameters. (**A-C**) Correlation analysis between FABP4 expression in tumor cells and various clinical parameters, including age (**A**), BMI (**B**), and tumor size (**C**). **(D-F)** Comparison of FABP4 expression levels in tumor cells of breast patients with different tumor grade (**D**), metastasis status (**E**), and patient vital status (**F**) (*p<0.05, ns: non-significant). (**G**) FABP4 expression across different breast cancer subtypes, including ER+/PR+/HER2-, ER-/PR-/HER2+, and ER-/PR-/HER2-(ns: non-significant). (**H, I**) Representative immunostaining for FABP4 in invasive frontline tumor cells and adjacent adipose tissue. No FABP4 was expressed in tumor cells (**H**) and FABP4 was expressed in tumor cells close to adipocytes (black arrows) (**I**). Scale bars: 100 μm. (**J**) Comparison of lymphovascular invasion percentage between patients with FABP4-positive (n=46) and patients with FABP4-negative (n=50) staining in front tumor cells (*p<0.05).

### FABP5 expression in tumor cells is associated with tumor metastasis

To further investigate the role of FABP5 in invasive breast cancer, we first analyzed the correlation between FABP5 expression in tumor cells and various clinical parameters, including age (Figure 5A), BMI (Figure 5B), tumor size (Figure 5C), tumor Elston-Ellis grade (Figure 5D), metastasis (Figure 5E) and patient vital status (Figure 5F). Similar to FABP4, FABP5 expression did not correlate with patient age, BMI and tumor size (Figure 5A-5C). However, FABP5 expression significantly corelated with tumor grade, tumor metastasis and patient vital status (Figure 5D-5F). Further analysis of FABP5 expression in tumor cells across different breast cancer subtypes demonstrated that both triple negative and HER2_subtypes exhibited significantly higher levels of FABP5 expression (p < 0.0001) compared to the hormone sensitive subtype (Figure 5G). Since triple negative and HER2+ subtypes are associated with a worse prognosis than the hormone sensitive subtype, our results suggest that FABP5 may serve as a marker for poor breast cancer prognosis. It is worth noting that in multiple specimens of tumor metastases, tumor cells within blood vessels in the primary tumor exhibited strong expression of FABP5 expression (Figure 5H-5J), further suggesting a possible pro-metastatic activity of FABP5 in tumor cells. Collectively, these findings indicate that intrinsic expression of FABP5 in breast cancer cells promotes metastatic spread, contributing to a worse prognosis in breast cancer patients.

**Fig. 5.**
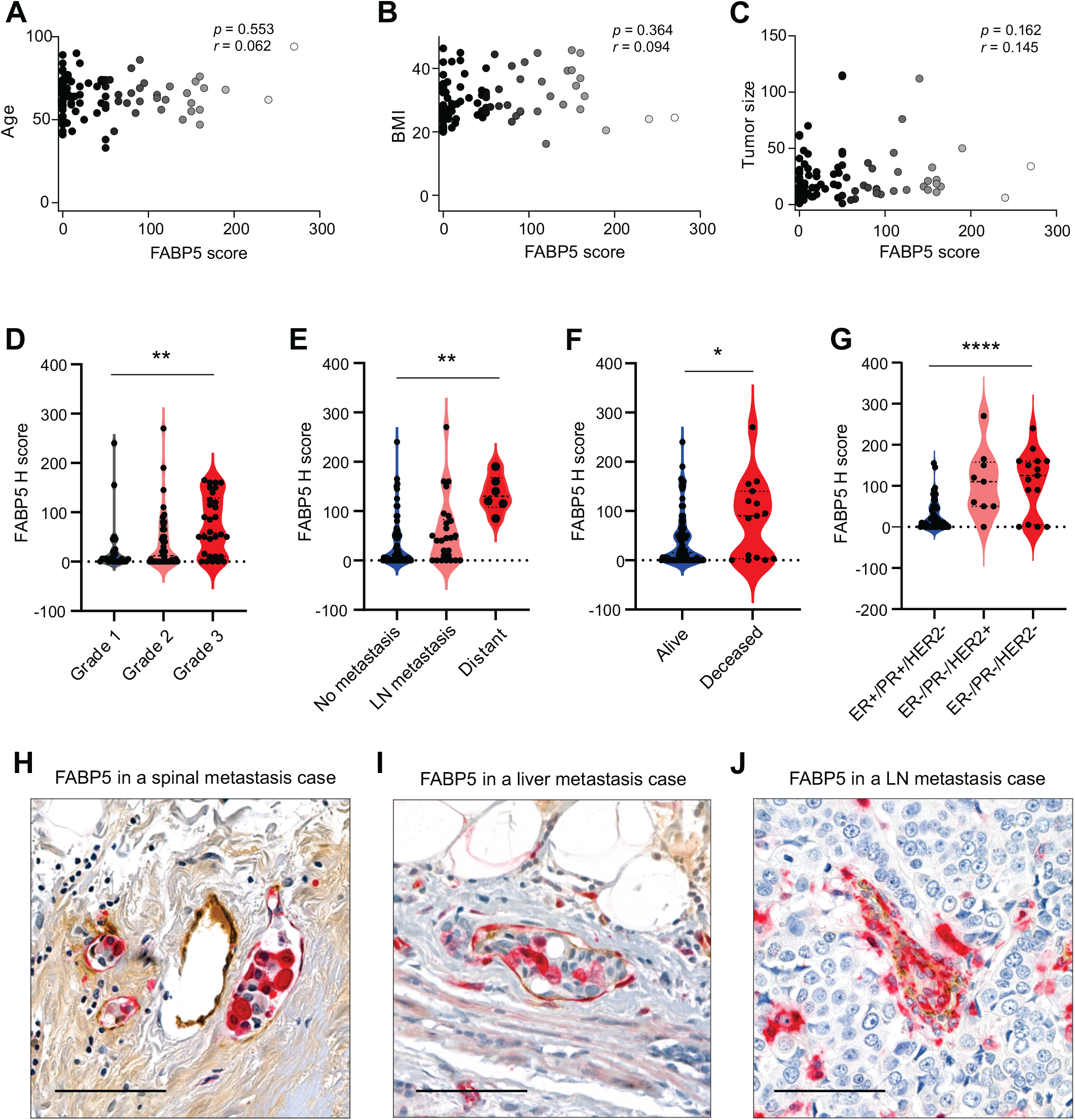
Correlations of FABP5 expression in tumor cells with clinical parameters. (**A-C**) Correlation analysis between FABP5 expression in tumor cells and various clinical parameters, including age (**A**), BMI (**B**), and tumor size (**C**). **(D-F)** Comparison of FABP5 expression levels in tumor cells of breast patients with different tumor grade (**D**), metastasis status (**E**), and patient vital status (**F**) (*p<0.01, **p<0.01). (**G**) FABP5 expression across different breast cancer subtypes, including ER+/PR+/HER2-, ER-/PR-/HER2+, and ER-/PR-/HER2-(****p<0.0001). (**H, I, J**) Representative images showing strong FABP5 expression in metastatic tumor cells within blood vessels in patient with spinal metastasis (**H**), liver metastasis (**I**) and lymph node (LN) metastasis (**J**).

## Conclusions

Breast epithelium, embedded in adipose tissue, is significantly influenced by lipid metabolism during both normal differentiation and carcinogenesis^21^. In breast cancer, fatty acid levels are generally higher than in normal tissue^22^. Cancer cells in the breast acquire lipids either from exogenous sources, such as surrounding adipose tissues or through their own *de novo* lipogenesis^23^. However, targeting lipids directly for therapeutic purposes is challenging because of the complexity of lipid species and the compensatory mechanisms within lipid synthesis pathways. Rather than focusing on lipids themselves, inhibiting the activity of FABPs, which act as lipid chaperones, may offer a novel strategy for treatment of lipophilic breast cancer.

In this study we demonstrate that FABP4 and FABP5 exhibit distinct expression patterns in breast tissue, with FABP4 predominantly expressed in stromal adipocytes, endothelial cells and macrophages, while FABP5 is primarily found in epithelial-derived tumor cells. Our findings present several key insights: 1) Epithelial-derived breast cancer cells typically do not express FABP4. However, invasive tumor cells near adipose tissue often express FABP4, suggesting that ectopic FABP4 may originate from neighboring adipocytes. This is supported by our observation of reduced adipocyte size at the tumor front and consistent with other animal model data^24^. Thus, FABP4 appears to facilitate the transport of exogenous fatty acids from the stroma to tumor cells, activating oncogenic signaling and enhancing lipid responses for breast cancer progression. 2) In aggressive breast cancers, especially in ER-negative subtypes, FABP5 is significantly upregulated in tumor cells compared to normal tissue or DCIS. Elevated FABP5 expression is positively correlated with tumor metastasis and decreased patient survival, suggesting that FABP5 functions as an intrinsic lipid chaperone within tumor cells, promoting lipid metabolism and tumor metastasis.

In summary, while FABP4 and FABP5 play distinct roles in regulating lipid availability for tumor cells, both are associated with poor prognoses in breast cancer. Given our recent development of multiple anti-FABP4 antibodies, which showed significant efficacy in reducing mammary tumor growth and metastasis in animal models^25^, FABP4 and FABP5 represent promising therapeutic targets, and blocking their activity may offer novel strategies for breast cancer treatment.

## Availability of data and materials

The datasets used and/or analyzed during the current study are available from the corresponding author on reasonable request.

## Abbreviations

FABP: Fatty acid binding protein
DCIS: Ductal carcinoma *in situ*
ER: Estrogen receptor
PR: Progesterone receptor
HER2: Human epidermal growth factor receptor 2
ALDH1: Aldehyde dehydrogenase 1
IHC: Immunohistochemical staining
vWF: von Willebrand factor
LN: Lymph node

## Acknowledgements

The authors would like to thank the Comparative Pathology Laboratory for histology services.

## Funding

National Institutes of Health grant R01AI137324 (BL), R01CA180986 (BL) and U01CA272424 (BL).

## Contributions

Conceptualization: B.L., E.D., Sample collection and Methodology: X.J., Y.Y., A.A, J.S., Z.W., X.C., J.H., S.L., M. C., Data analysis: X.J., B.L., Y.X., Writing—original draft: X.J., B.L., Writing—review and editing: S.S., E.D., Supervision: B.L. Funding acquisition: B.L., S.S. All authors read and approved the final manuscript.

## Ethics declarations

Ethics approval and consent to participate. All experiments were performed according to the approval by the institutional review board at the University of Iowa, Iowa City, IA.

## Competing interests

The authors declare no competing interests.

## Notes

### Competing Interest Statement

The authors have declared no competing interest.

